# Transferrin-induced signaling through transferrin receptor and AKT kinase mediates formation of Rab8- and MICAL-L1-positive tubules involved in receptor recycling

**DOI:** 10.1101/2023.02.06.527405

**Authors:** Nandini Rangaraj, Vipul Vaibhava, Cherukuri Sudhakar, Shivranjani C Moharir, Ghanshyam Swarup

**Affiliations:** CSIR-Centre for Cellular and Molecular Biology, Hyderabad-500007, India

**Keywords:** Transferrin, transferrin receptor recycling, RAB8, MICAL-L1, AKT, TBC1D17, endocytosis

## Abstract

Transferrin and its receptor play an important role in iron homeostasis. Binding of transferrin to its receptor (TFRC, transferrin receptor protein 1) initiates endocytic trafficking and subsequent recycling of TFRC to the plasma membrane. RAB8-positive tubules emanating from the endocytic recycling compartment play an important role in receptor recycling. However, the signaling pathways or mechanisms that mediate formation of RAB8-positive tubules are not clear. Here, we have investigated the role of transferrin-induced signaling in the regulation of RAB8- and MICAL-L1-positive tubules. Addition of transferrin to the serum starved HeLa cells resulted in enhanced recruitment of RAB8 as well as MICAL-L1 to the tubules, which was mediated by TFRC. Dynasore, an inhibitor of dynamin and endocytosis, completely blocked transferrin-induced formation of RAB8/MICAL-L1-positive tubules. RAB8 showed strong colocalization with MICAL-L1 on the tubules. Blocking of SRC or AKT kinase activity by specific inhibitors abolished transferrin-induced recruitment of RAB8 and MICAL-L1 to the tubules. Recycling of transferrin receptor was inhibited by blocking of AKT activity. TBC1D17, a GTPase activating protein for RAB8, inhibited RAB8/MICAL-L1-positive tubule formation. A phospho-mimicking mutant S366D of TBC1D17 did not inhibit formation of RAB8-positive tubules. Overall, these results show that transferrin induces TFRC mediated signaling dependent on endocytosis that is essential for the formation of RAB8- and MICAL-L1-positive tubules involved in recycling of transferrin receptor. Our results also show that AKT regulates transferrin-induced formation of RAB8- and MICAL-L1-positive tubules, which might be mediated by phosphorylation of TBC1D17.

## Introduction

Endocytosis is an essential cellular process that is required for uptake of nutrients from the extracellular environment and delivery of membrane components (lipids, proteins and their bound ligands) to various intracellular locations. Generally, endocytosis is receptor mediated and involves identification of nutrient/signal by the receptor at the plasma membrane, its internalization and release inside the cell [1]. The receptor can be recycled back to the plasma membrane for further cycles to be performed or transported to lysosomes for degradation. Transferrin (Tfn), a glycoprotein present in the blood plasma, is involved in iron transport from outside the cell through receptor mediated endocytosis [2. 3]. Iron-bound transferrin binds to its receptor TFRC (transferrin receptor protein 1) to initiate endocytosis leading to formation of vesicles, which move to the early endosomes [4]. Iron is released inside the endosomes due to acidic environment and transported to the cytoplasm through transport proteins [2-5]. The transferrin-TFRC complex is recycled back to the plasma membrane from early endosomes (fast recycling) or moves to the recycling endosomes. The Tfn-TFRC complex is also recycled to the plasma membrane from the recycling endosomes (slow recycling). Tfn has two receptors TFR1 (TFRC, gene *TFRC*) and TFR2 [4, 6]. The TFRC is expressed in almost all the cells. While most of the TFRC is recycled back to the plasma membrane, a small fraction is constitutively degraded in lysosomes through an autophagic pathway that is dependent on RAB12 [7, 8]. TFRC degradation in lysosomes also takes place under conditions of iron overload through an unconventional endocytic pathway [9]. The mechanisms that control recycling of Tfn-TFRC are not completely understood.

RAB GTPases control most of the steps of vesicle trafficking pathways including endocytic trafficking and recycling. Several RABs such as RAB4, 5, 8, 11, 22 and 35 are involved in regulating one or more steps of transferrin trafficking and recycling [10-15]. RAB8 GTPase is involved in mediating Tfn trafficking and recycling from the recycling endosomes [14, 15]. It is required for transport of Tfn containing vesicles from early endosomes to the recycling endosomes [14]. It has been shown to localize to recycling tubular endosomes (RTEs) as seen by its colocalization with MICAL-L1 and EHD1 proteins. RTEs are necessary for the delivery of membrane and proteins to the plasma membrane. Activated form of RAB8 (RAB8-GTP) directly interacts with its effector, optineurin (OPTN), which in turn forms a complex with TFRC [15, 16]. Activated form of RAB8 is recruited to the tubules originating from the endocytic recycling compartment. Recruitment of RAB8 to these tubules is involved in mediating recycling of TFRC to the plasma membrane. The activity of RAB8 is regulated by a GTPase activating protein (GAP), TBC1D17, which indirectly interacts with RAB8 through an adaptor protein Optineurin [15]. In addition to RAB8, these tubules contain MICAL-L1 and EHD1 proteins [17, 18]. MICAL-L1 is required for recycling of TFRC and beta1 integrin [17, 18]. The interaction of MICAL-L1 with RAB8 facilitates an indirect interaction between EHD1 and RAB8 [18]. MICAL-L1 is considered as an effector of RAB8 because it interacts with the activated form of RAB8.

Although the primary function of transferrin is in the process of iron uptake, it is also involved in regulation of cell growth and myelination of axons in the central nervous system [3, 19-28]. Signal transduction induced by binding of transferrin to its receptor leads to activation of several signaling proteins such as SRC, FYN, AKT, ERK-1/ERK-2 kinases {3, 29-31]. Tfn induced signaling leads to phosphorylation of cortactin and dynamin by the SRC kinase, which gets activated [29, 30]. These signaling processes induced by Tfn (Src kinase activation and phosphorylation of cortactin and dynamin by the SRC kinase) are required for endocytosis of transferrin [30]. Phosphorylation of TFRC on tyrosine has been observed under certain conditions [32]. However, the role of Tfn-induced signaling in the control of Tfn/TFRC trafficking and recycling is not clear.

Here, we have examined the role of Tfn-induced signaling in the formation of RAB8- and MICAL-L1-positive tubules that are involved in recycling of Tfn/TFRC. Our results show that Tfn treatment of the serum-starved HeLa cells induces formation of RAB8- and MICAL-L1-positive tubules dependent on endocytosis. AKT kinase plays an essential role in the formation of these tubules and in the recycling of transferrin. We have also explored the role of TBC1D17, a RAB8 GTPase activating protein, and its possible phosphorylation by AKT in transferrin-induced formation of the RAB8-positive tubules.

## Materials and Methods

### Reagents

Mouse monoclonal anti-RAB8 antibody was from BD Biosciences (#610845) (San Jose, CA, USA); rabbit polyclonal anti-RAB8 antibody was from Sigma (#R5530); mouse polyclonal anti-MICAL-L1 antibody was from Novus Biologicals (H00085377-B01P); rabbit polyclonal anti-Akt antibody was from Cell signalling (#9272); mouse monoclonal anti-transferrin receptor antibody was from Calbiochem (#GRO9L); anti-mouse and anti-rabbit Cy3 conjugated secondary antibodies were from Amersham; anti-rabbit and anti-mouse Alexafluor-488 conjugated secondary antibodies, Alexa Fluor 546-labelled transferrin (#T23364) were from Molecular probes; Akt inhibitor AI4 was from Calbiochem (#124011); Src family kinase inhibitor SU6656 was from Sigma (##9692); Wortmannin (phosphatidylinositol 3-kinase inhibitor) was from Calbiochem (#681675); bovine serum albumin (BSA) (# A9418) and unlabelled human holo-transferrin (#T0665) were from Sigma.

### Expression plasmids

Plasmid expressing dominant negative AKT was kindly provided by Dr Gopal Kundu of National Centre for Cell Science, Pune, India. TBC1D17 and its R381A mutants with GFP and HA tags have been described earlier [15]. S366D mutant of TBC1D17 was made by site-directed mutagenesis [15].

### Cell culture and transfection

HeLa cells were grown as monolayers in T-25 flasks, in a humidified atmosphere at 5% CO_2_ and 37°C in DMEM (Dulbecco’s Modified Eagle’s medium) containing 10% fetal calf serum. Experiments were carried out by plating cells on coverslips. Transfection was done using Lipofectamine™ 2000 (Invitrogen Life technologies, CA, USA).

### Indirect immunofluorescence and Confocal Microscopy

For staining of endogenous RAB8 and MICAL-L1, cells were fixed with 3.7% formaldehyde for 10min, permeabilized for 6min with 0.5% Triton X-100 and 0.05% Tween 20 (Sigma) in PBS. Permeabilized cells were incubated in blocking solution (2% BSA in PBS) for 1hr at room temperature. The cells were then incubated with RAB8 antibody (1:200 dilution) in blocking solution for 2hrs at room temperature followed by 10-12hrs at 4°C. Cells were then washed and incubated with Cy3 conjugated anti-rabbit antibody or anti-mouse antibody as the case may be (1:1200 dilution) for 1hr at room temperature. For dual labelling of RAB8 and MICAL-L1, rabbit polyclonal RAB8 antibody was used. For staining of endogenous MICAL-lL1, after washing the cells were incubated with MICAL-L1 antibody (1:200 dilution) in blocking solution for 3hrs at room temperature. Cells were then washed and incubated with Alexa fluor-488 conjugated anti-mouse antibody (1:400 dilution) for 1hr at room temperature. Cells were then washed with PBS and mounted with mountant containing antifade and DAPI.

For staining of overexpressed dominant negative AKT, after fixing, permeabilizing and blocking the cells were incubated with AKT antibody (1:200 dilution) in blocking solution for 2hrs at room temperature. The cells were then washed and incubated with Alexafluor-488 conjugated anti-rabbit antibody (1:400 dilution) for 1hr at room temperature. Cells were washed and labelled for endogenous RAB8 or MICAL-L1 as mentioned above.

Labelled cells were observed using LSM 510 NLO Confocal Microscope (Carl Zeiss Micro imaging, Jena, Germany) or TCS SP5 Leica Confocal Microscope (Leica Micro systems, Mannheim, Germany). For imaging Alexa-488 and Cy3, a 488nm argon laser and 561nm DPSS (or 543nm HeNe) lasers were used respectively. Serial optical sections in the Z-axis of the cells were collected at 0.35μm intervals with a 63X oil immersion objective lens (NA 1.4). Generally, 2-4 optical sections were projected using LSM510 (version 3.2) or the Leica LAS AF software. Images were further processed using Adobe Photoshop for making figures as described earlier [15].

### Analysis of RAB8- and MICAL-L1-positive tubules

HeLa cells induced with transferrin or vehicle alone, and cells after transfection with required plasmid or untransfected cells after transferrin/BSA induction were labelled for RAB8 and/or AKT and MICAL-L1. Images were acquired on Confocal microscope and projected using the software. Cells containing at least one distinct tubule per cell with length greater than 10μm were counted. At least 100 cells in case of induced/untransfected cells, and 40 cells in case of transfected cells were counted for each set of experiment.

### Treatment of cells

For time course experiments, HeLa cells grown on coverslips were washed and pre-incubated with serum free DMEM for 2hrs. These cells were then incubated with 10μg/ml of unlabelled transferrin in serum free medium for 1hr at 4°C. Bovine serum albumin (BSA) at a concentration of 10μg/ml in serum free DMEM was used as a control for this experiment. The cells were then shifted to 37°C to allow uptake of transferrin for different time intervals (i.e. 0, 10, 15, 20, 30min). The cells were removed from 37°C incubator at the specified time intervals, washed with PBS twice and fixed with 3.7% formaldehyde. Later the cells were labelled for endogenous RAB8 and MICAL-L1 as described.

To observe the effect of AKT inhibitor, the cells were serum starved for 2hrs and incubated with 5μM of AKT inhibitor AI4 in dimethyl sulfoxide (DMSO) along with 10μg/ml of unlabelled transferrin in serum free medium for 1hr at 4°C. DMSO at 0.05% and BSA at 10μg/ml were used as controls. The cells were then shifted to 37°C to allow uptake of transferrin for 20 minutes. The cells were then washed, fixed and labelled for endogenous RAB8 and MICAL-L1.

To observe the effect of over-expressed dominant negative AKT, the cells were transfected with dominant negative AKT expression plasmid (400ng), and 22-24 hrs after transfection, the cells were pre-incubated with serum free medium for 2hrs. These cells were then incubated with 10μg/ml of unlabelled transferrin for 1hr at 4°C in serum free medium. BSA at 10μg/ml was used as control. The cells were then shifted to 37°C for 20minutes, washed, fixed and immuno-stained for AKT followed by endogenous MICAL-L1 or RAB8.

To know whether transferrin-induced tubule formation was mediated through transferrin receptor, HeLa cells grown on coverslips were kept in serum free DMEM for 2hrs and later the cells were pre-incubated with 5μg/ml of TFRC antibody or control IgG in serum free medium for 30min at 37°C and incubated along with 10μg/ml of unlabelled transferrin or BSA for 1hr at 4°C.The cells were then shifted to 37°C to allow uptake of transferrin for 20min. The cells were then washed, fixed and labelled for endogenous RAB8 and MICAL-L1.

To explore the role of endocytosis in transferrin-induced tubule formation, HeLa cells grown on coverslips were kept in serum free DMEM for 2hrs.The cells were then pre-incubated with 80μM Dynasore [33] or 0.1% of solvent, DMSO for 30min at 37°C and then incubated along with 10μg/ml of unlabelled transferrin or BSA for 1hr at 4°C. The cells were then shifted to 37°C to allow uptake of transferrin for 20min.

To explore the role of kinases in transferrin-induced tubule formation, HeLa cells were serum starved for 2hrs and then incubated with 10μM of the SRC-inhibitor SU6656 or 250nM of the phosphatidylinositol 3-kinase (PI3K) inhibitor along with 10μg/ml of transferrin or BSA for 1hr at 4°C. DMSO at 0.1% was used as the solvent control. Then the cells were shifted to 37°C for 20min to allow uptake of transferrin. Cells were then washed, fixed and labelled for RAB8 and MICAL-L1.

### Statistical Analysis

Graphs represent average ±SD values. Statistical significance was calculated using Students T-test. When significant values were observed, P-values for pair wise comparison were calculated by using two-tailed T-test. P-values less than 0.05 were considered significant.

## Results

### Transferrin induces formation of RAB8- and MICAL-L1-positive tubules

Tubular structures originating from the endocytic recycling compartment (ERC) are involved in the recycling of receptor proteins to the plasma membrane. Activated form of RAB8 preferentially associates with these tubules, which also contain MICAL-L1 and EHD1 proteins [14, 15, 18]. However, the mechanisms that regulate formation of these RAB8-positive tubules are not clear. Since these RAB8-positive tubules are involved in mediating Tfn-TFRC recycling, we hypothesized that transferrin-induced signaling may be involved in regulating the formation of these tubules. Therefore, we examined the effect of transferrin (holo-transferrin) treatment of the cells on the formation of RAB8-positive tubules. We carried out this experiment under the conditions frequently used for transferrin uptake assays. HeLa cells were serum starved for 2 hours, incubated with unlabelled transferrin at 4°C for one hour and then the incubation was continued at 37°C for 5-30 min. The cells were then fixed, stained with RAB8 and MICAL-L1 antibodies, and examined by confocal microscopy (Fig. 1A). Cells treated with bovine serum albumin were used as controls. Treatment of cells with transferrin resulted in a time dependent increase in the number of cells showing RAB8-positive tubules (Fig. 1A, B). Maximum increase in the percentage of cells showing RAB8-positive tubules was seen after 20 minutes of treatment with transferrin at 37°C. Treatment of cells with transferrin also resulted in an increase in the number of cells showing MICAL-L1-positive tubules (Fig. 1C). Time course of transferrin-induced increase in MICAL-L1 tubules was similar to that of RAB8 tubules (Fig. 1B, C). There was strong colocalization of RAB8 with MICAL-L1 on the tubules (Fig. 1D). The number of cells showing MICAL-L1-positive tubules was always more than the number of cells showing RAB8-positive tubules. Almost all the cells showing RAB8-positive tubules showed MICAL-L1-positive tubules, but some of the cells showing MICAL-L1 positive tubules did not show RAB8-positive tubules.

**Figure 1:**
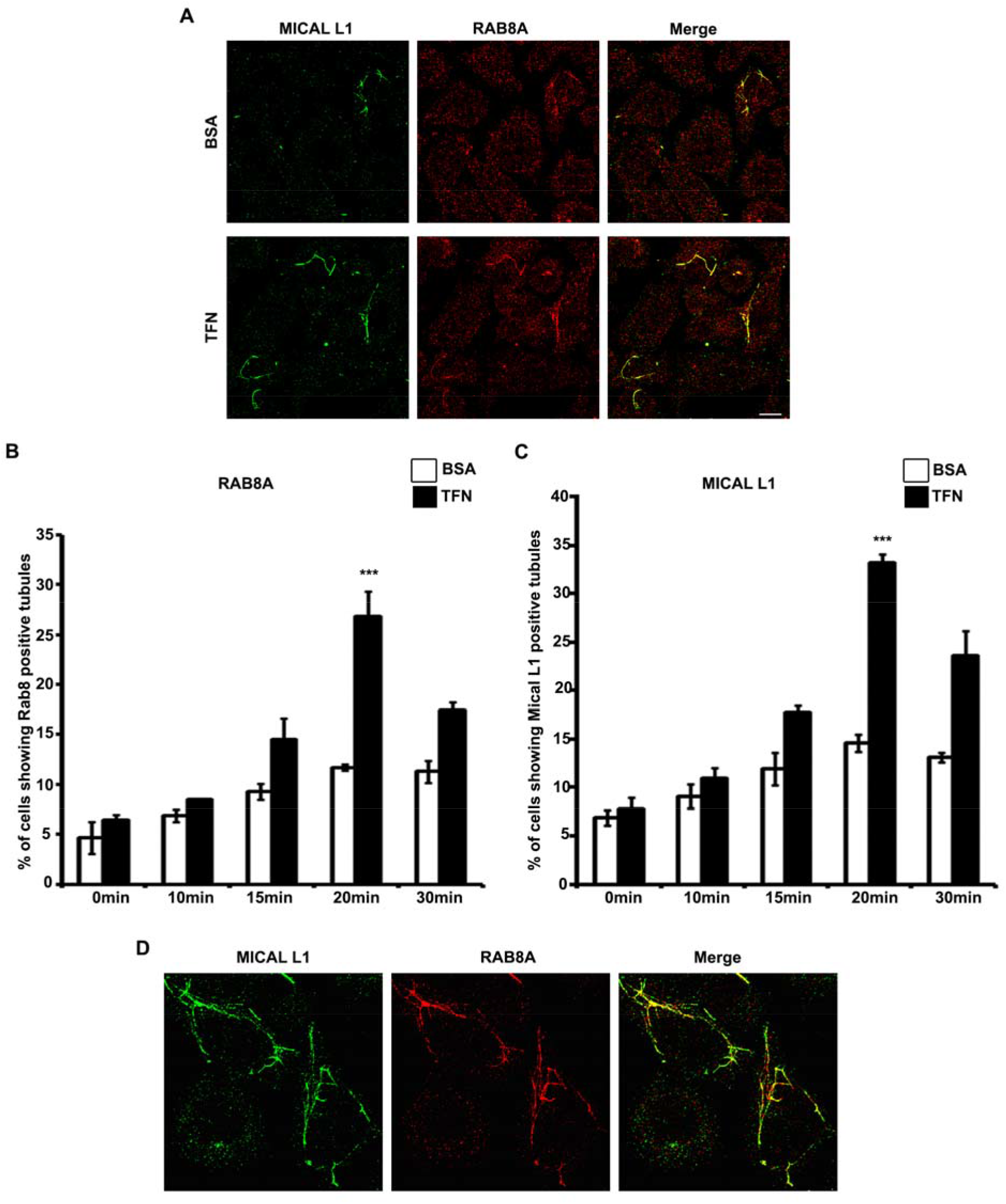
Transferrin induces formation of RAB8 and MICAL-L1-positive tubules. Hela cells grown on coverslips were serum starved for 2 hours, treated either with 10μg/ml of transferrin (TFN)) or BSA, kept at 4°C for one hour and incubated at 37°C for indicated time. The cells were then fixed, stained for endogenous Rab8 and MICAL-L1 and examined by confocal microscope. Representative confocal images after 20 minutes of treatment with BSA and transferrin are shown (A). Bar diagrams show quantitation of the percentage of cells exhibiting RAB8 tubules (B), or MICAL-L1 tubules (C) at indicated time. Data represent mean ±s.d. from 3 independent experiments done with 265-332 cells. High resolution images show colocalization of Rab8 tubules with MICAL-L1 tubules (D). Scale bar, 10 μm.

### Transferrin-induced formation of RAB8- and MICAL-L1 positive tubules is mediated by transferrin receptor

Transferrin receptor TFRC is expressed in almost all the cells. We examined the role of TFRC in transferrin-induced formation of RAB8 and MICAL-L1 tubules by using a monoclonal antibody that is known to prevent binding of transferrin to TFRC. HeLa cells grown on coverslips were serum starved for 2 hours, pretreated with TFRC antibody or control antibody for 30 min at 37°C and then incubated with transferrin or BSA (control) for 1 hour at 4°C. The cells were then shifted to 37°C for 20 min. Pretreatment with TFRC antibody resulted in significant reduction in the number of cells showing transferrin-induced formation of RAB8 as well as MICAL-L1-positive tubules (Fig. 2A, B, C). There was no significant change in the percentage of cells showing basal level of RAB8 or MICAL-L1 positive tubules upon pretreatment with TFRC antibody (Fig. 2A, B, C). These results suggest that transferrin-induced formation of RAB8 and MICAL-L1 positive tubules is mediated by TFRC.

**Figure 2:**
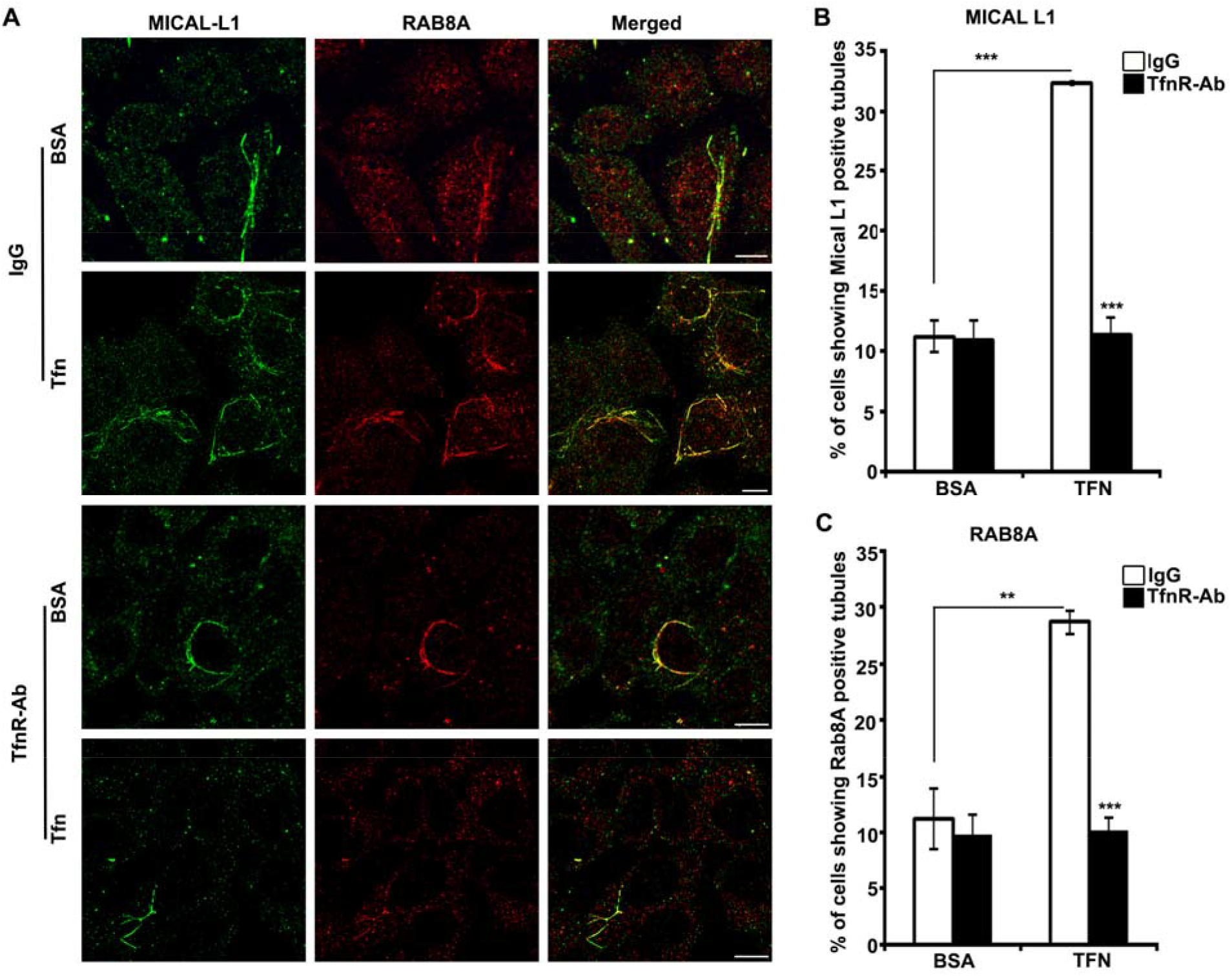
Transferrin-induced formation of RAB8 and MICAL-L1 positive tubules is mediated by TFRC. Hela cells grown on coverslips were serum starved for 2 hours, preincubated with TFRC blocking antibody or control IgG for 30 minutes and then treated with either transferrin or BSA, kept at 4°C for 1 hour followed by incubation at 37°C for 20 minutes. The cells were then fixed, stained for endogenous RAB8 and MICAL-L1 and examined by confocal microscope. Representative confocal images are shown (A). Bar diagrams show quantitation of the percentage of cells exhibiting MICAL-L1 tubules (B), or RAB8 tubules (C). Data represent mean± s.d. from 3 experiments done with 298-349 cells. *** P<0.0001, ** P<0.01. Scale bar, 10 μm.

### Endocytosis is required for transferrin-induced formation of RAB8 and MICAL-L1-positive tubules

Signaling by transmembrane receptors sometimes requires endocytosis of the receptor. Since TFRC is endocytosed readily, we examined the requirement of endocytosis for transferrin-induced formation of RAB8 and MICAL-L1-positive tubules. For this purpose, we used Dynasore, a small molecule and a potent inhibitor of dynamin GTPase (and therefore, inhibits dynamin-dependent endocytosis) that is known to block the uptake, trafficking and intracellular accumulation of transferrin [33]. Pretreatment of cells with Dynasore reduced transferrin-induced increase in RAB8 and MICAL-L1-positive tubules (Fig. 3A, B, C) whereas basal level of tubule formation was not affected significantly. These results suggest that endocytosis is required for transferrin-induced formation of RAB8 and MICAL-L1 positive tubules.

**Figure 3:**
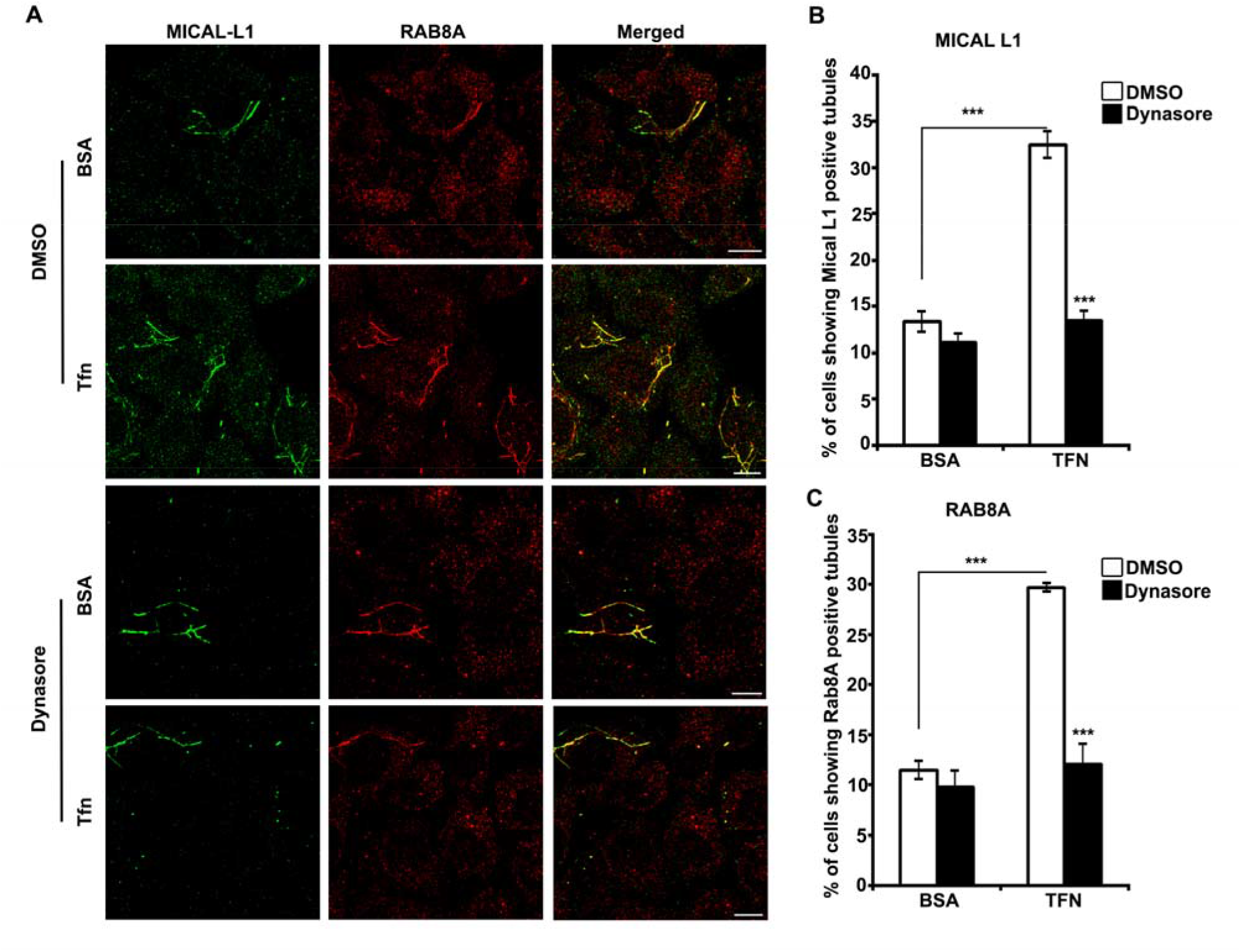
Endocytosis is required for transferrin-induced formation of RAB8 and MICAL-L1 positive tubules. Hela cells grown on coverslips were serum starved for 2 hours, preincubated with Dynasore (80μM) or DMSO (0.1%) for 30 minutes and then treated with either transferrin or BSA, kept at 4°C for 1 hour followed by incubation at 37°C for 20 minutes. The cells were then fixed, stained for endogenous RAB8 and MICAL-L1, and examined by confocal microscope. Representative confocal images are shown (A) Scale bar: 10μm. Bar diagrams show quantitation of the percentage of cells exhibiting MICAL-L1 tubules (B), or RAB8 tubules (C). Data represent mean± s.d. from 3 experiments done with 328-347 cells. *** P<0.0001, ** P<0.01.

### AKT activity is required for the transferrin-induced formation of RAB8 and MICAL-L1 positive tubules

AKT is a protein kinase involved in regulating many cellular functions. Activation of AKT has been reported upon treatment of oligodendroglial precursor cells with transferrin [3]. We examined the role of AKT in transferrin-induced formation of RAB8- and MICAL-L1-positive tubules by using a chemical inhibitor and also by using a kinase-dead mutant of AKT. Treatment of cells with AKT inhibitor resulted in complete inhibition of transferrin-induced increase in the number of cells showing MICAL-L1-positive tubules whereas cells treated with the solvent (DMSO) showed transferrin-induced increase in number of cells with MICAL-L1-positive tubules (Fig. 4A, B). However, there was only a small effect of AKT inhibitor on basal level of MICAL-L1-positive tubules. Similarly, treatment of cells with AKT inhibitor but not with solvent resulted in complete loss of transferrin-induced formation of RAB8-positive tubules (Fig. 4A, C). These results show that AKT activity is required for transferrin-induced formation of RAB8- and MICAL-L1-positive tubules. To provide further evidence for the role of AKT in transferrin-induced RAB8-positive tubule formation, we used a kinase-dead mutant of AKT. HeLa cells were transfected with a plasmid expressing kinase-dead AKT and, after 24 hours treated with transferrin. These cells were stained for AKT and MICAL-L1 using specific antibodies and the cells were analyzed by confocal microscopy. Cells expressing kinase-dead AKT showed complete inhibition of transferrin-induced increase in MICAL-L1-positive tubules (Fig. 4D, E). Even in cells not treated with transferrin, there was some decrease in the number of cells showing MICAL-L1-positive tubules upon expression of kinase-dead AKT. In similar experiments, expression of kinase-dead AKT resulted in complete inhibition of transferrin-induced increase in the number of cells showing RAB8-positive tubules (Fig. 4F, G). These results show that AKT mediates transferrin-induced formation of MICAL-L1-positive and RAB8-positive tubules.

**Figure 4:**
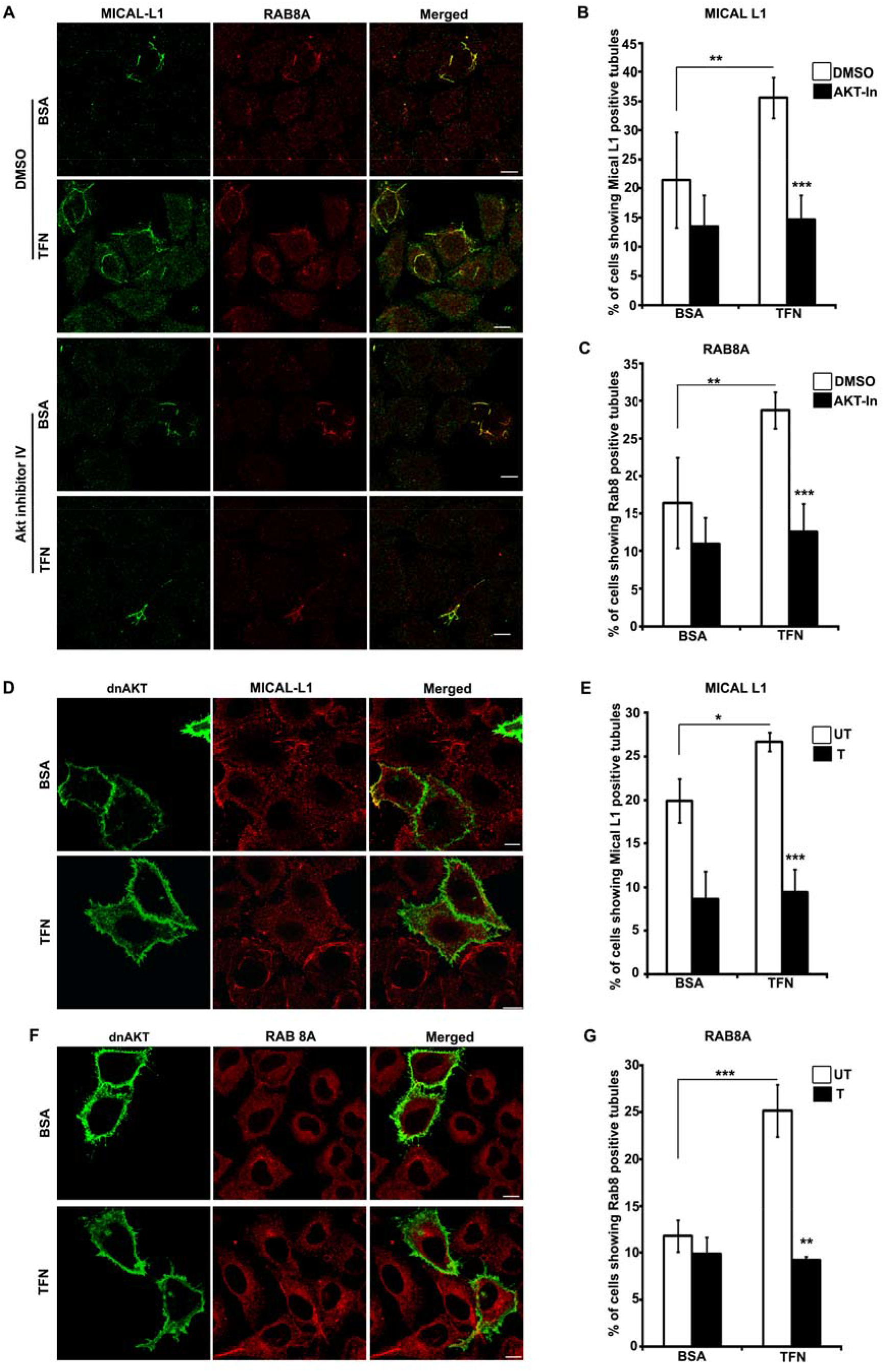
AKT activity is required for transferrin-induced formation of RAB8 and MICAL-L1 positive tubules. Hela cells grown on coverslips were serum starved for 2 hours, and treated with either BSA or transferrin and with DMSO (0.05%) or Akt inhibitor AI-4, kept at 4°C for 1 hour followed by incubation at 37°C for 20 minutes. Cells were fixed and stained for endogenous RAB8 and MICAL-L1 and observed by confocal microscope. Representative confocal images are shown (A) Scale bar: 10 μm. Bar diagrams show quantitation of the percentage of cells exhibiting MICAL-L1-positive (B) or RAB8-positive (C) tubules in each scoring category. Data from three separate experiments done with 411-503 cells are shown as the mean± s.d.***P<0.0001, *P<0.05. (D, E, F) Hela cells were transfected with dominant negative AKT. After 22 hours the cells were serum starved for 2 hours, treated with either BSA or transferrin, kept at 4°C for 1 hour followed by incubation at 37°C for 20 minutes. Cells were fixed and stained for endogenous RAB8 or MICAL-L1 and observed by confocal microscope. Representative confocal images are shown (D, F). Scale bar: 10 μm. Bar diagrams show percentage of cells exhibiting MICAL-L1 positive tubules (E) (transfected (T): 154cells; untransfected (UT): 602-608cells) or RAB8-positive tubules (G) (transfected: 99-119cells; untransfected: 360-443cells) in each scoring category. Data from three separate experiments are shown as the mean±s.d., ***P<0.0001, **P<0.01.

### AKT activity is required for transferrin receptor recycling

RAB8-positive tubules are involved in mediating transferrin receptor recycling from the endocytic recycling compartment (ERC) to the plasma membrane. Since AKT is required for transferrin-induced RAB8-positive tubule formation, we examined the role of AKT in transferrin receptor recycling. HeLa cells were transfected with the plasmid expressing kinase-dead AKT and transferrin uptake and recycling assays were carried out by incubating the cells with Alexa546-labelled transferrin for 20 minutes, and for recycling assay the cells were then washed with complete medium for 45-minutes at 37°C (chase). The cells were fixed either before or after the chase and examined by confocal microscopy. Expression of kinase-dead mutant of AKT resulted in significantly reduced uptake of transferrin as determined by quantitative analysis (Fig. 5A, B). GFP expressing cells were used as control, which did not show any inhibition of transferrin uptake. The level of TFRC was not altered in the cells expressing kinase-dead mutant of AKT, as seen by the western blot (Fig. 5C). After the chase, more labelled transferrin was seen in the cells expressing kinase-dead AKT, as compared with non-expressing cells or GFP expressing cells (Fig. 5D, E). These results suggest that AKT activity is required for transferrin receptor recycling.

**Figure 5:**
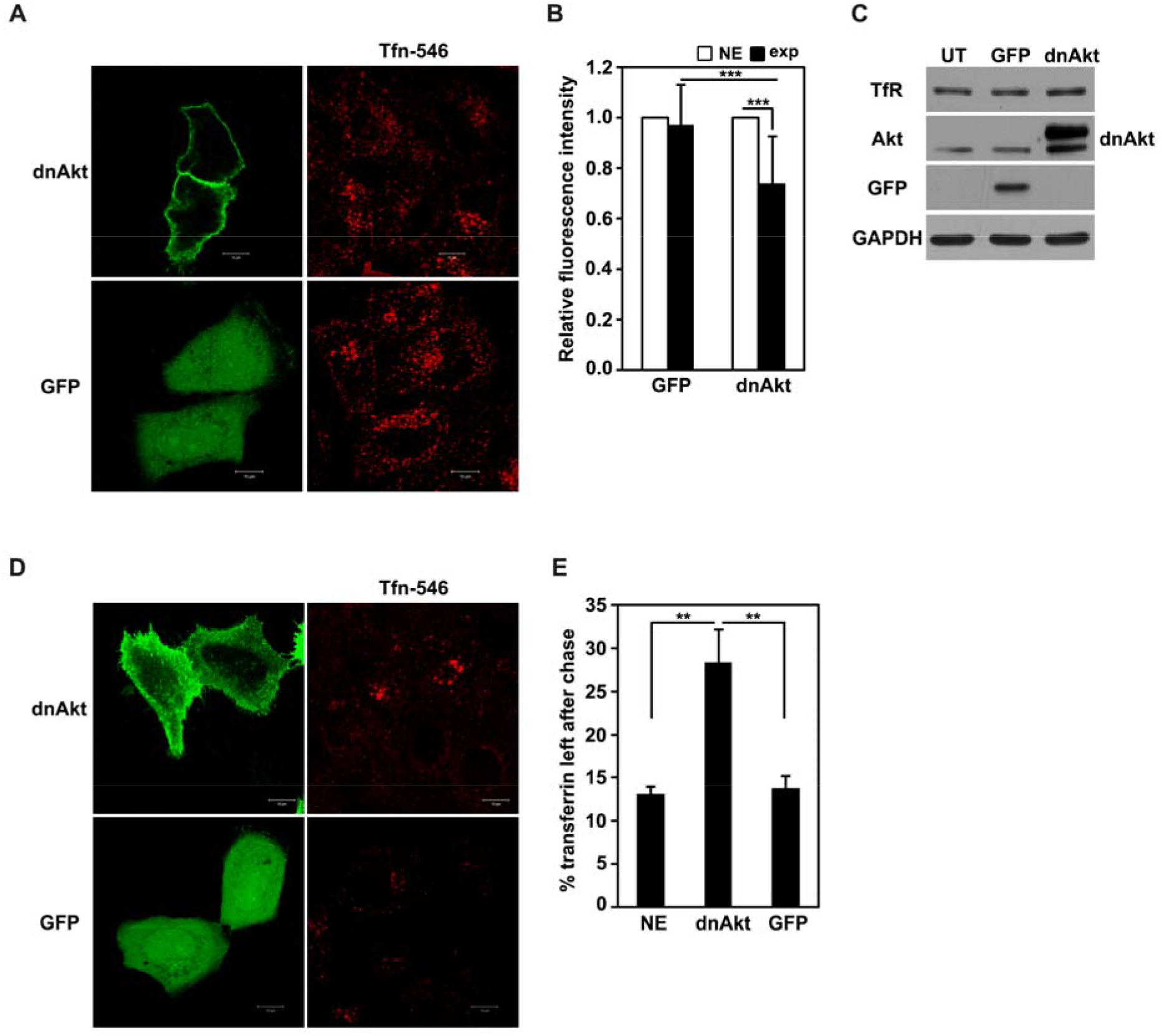
AKT activity is required for recycling of transferrin receptor. (A) Hela cells were transfected with dominant negative AKT or GFP. After 22 hours, a transferrin uptake assay was performed using Alexa-Fluor-546-conjugated transferrin (Tfn546). (B) Graph shows the relative fluorescence intensity of endocytosed transferrin in cells expressing dnAkt or GFP. ***P<0.001. (C) HeLa cells were transfected with GFP or dnAkt. After 24 hours, lysates were made, separated by SDS-PAGE and analysed by western blotting (IB) using anti-GFP, anti-Akt, anti-TFRC and anti-GAPDH antibodies. UT, untransfected cells. (D) HeLa cells grown on coverslips were transfected with GFP or dnAkt. After 24 hours, the cells were serum starved for 2 hours, incubated with Alexa-546-labelled transferrin (Tfn-546) for 20 minutes and then fixed, or washed twice with PBS and incubated in complete medium for 45 minutes (chase). The fixed cells were examined by confocal microscopy. (E) Quantitative analysis was carried out to calculate the percentage of transferrin remaining after the chase in expressing and non-expressing (NE) cells. **P<0.01. Scale bar: 10 μm.

### TBC1D17 inhibits formation of RAB8- and MICAL-L1-positive tubules

GTPase activating proteins (GAPs) with TBC domain regulate the activity of RAB GTPases [34, 35]. TBC1D17 is a GAP for RAB8 GTPase. It inhibits some of the RAB8 mediated functions, such as formation of RAB8-positive tubules and recycling of TFRC [15]. MICAL-L1 is an effector for RAB8, which crosslinks RAB8 and EHD1 [18]. We analyzed the effect of TBC1D17 on MICAL-L1 and RAB8-positive tubules. HeLa cells were transfected with GFP-TBC1D17 or its catalytically inactive R381A mutant, stained for MICAL-L1 and observed by confocal microscopy. TBC1D17 expressing cells showed a significant reduction in the formation of MICAL-L1-positive tubules as compared to untransfected cells or control GFP expressing cells or R381A expressing cells (Fig. 6 A, B). These results indicate towards a role of RAB8 in the recruitment of MICAL-L1 on the tubules. We also examined the role of TBC1D17 in RAB8-positive tubule formation. HeLa cells expressing GFP-TBC1D17 or its catalytically inactive R381A mutant were fixed and stained for RAB8 and observed by confocal microscopy. Expression of TBC1D17 inhibited recruitment of RAB8 on the tubules (Fig. 7A, B, C). However, expression of catalytically inactive R381A mutant of TBC1D17 resulted in a significant increase in the formation of RAB8-positive tubules (Fig. 7A, B, C).

**Figure 6:**
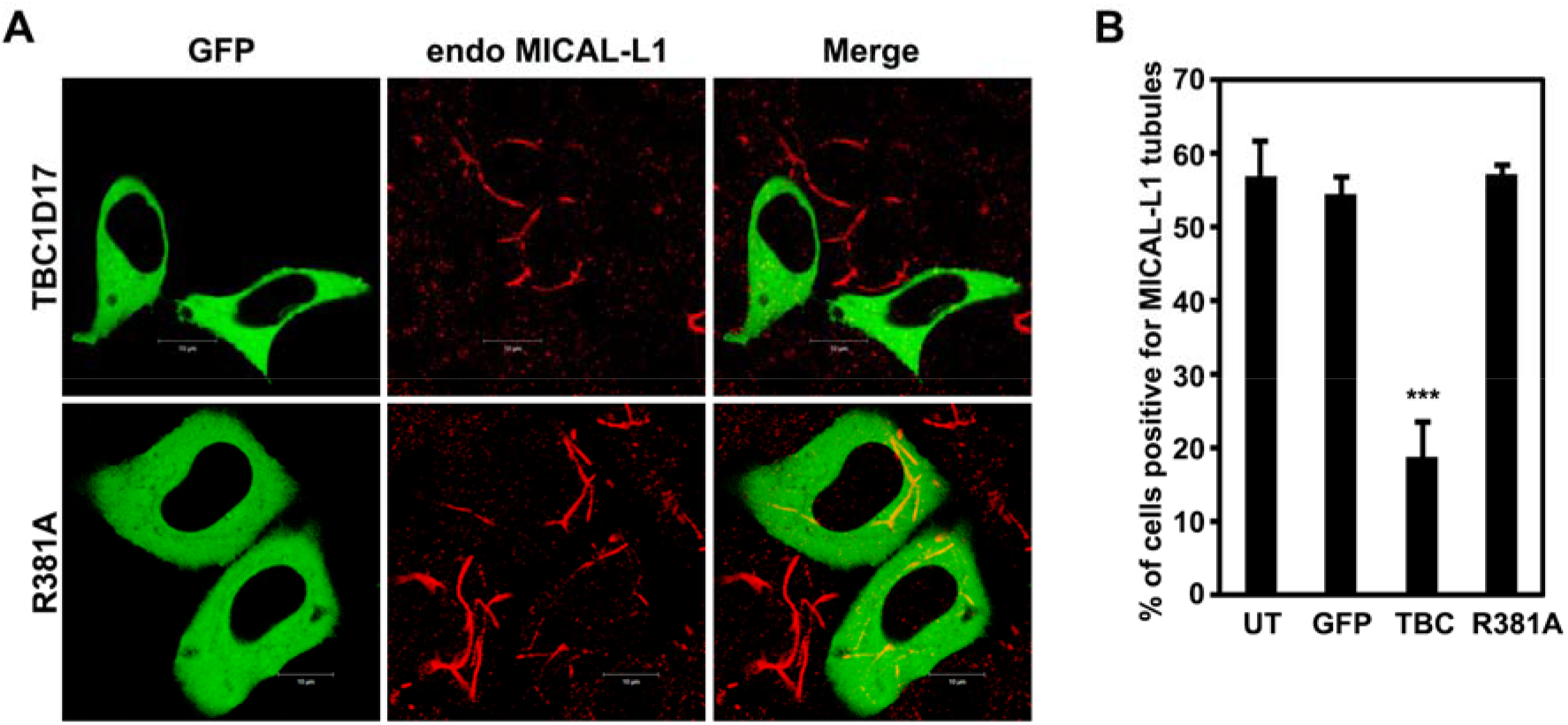
TBC1D17 inhibits formation of MICAL-L1 tubules. (A) Hela cells transfected with TBC1D17 or its catalytically inactive R381A mutant were stained for endogenous MICAL-L1. Confocal microscopy revealed that expression of TBC1D17 led to reduction in cells showing MICAL-L1 tubules. Scale bar: 10 μm. (B) The graph shows the percentage of cells exhibiting MICAL-L1-positive tubules in each scoring category. Data from three separate experiments are shown as the mean ± s.d. ***P<0.001.

**Figure 7:**
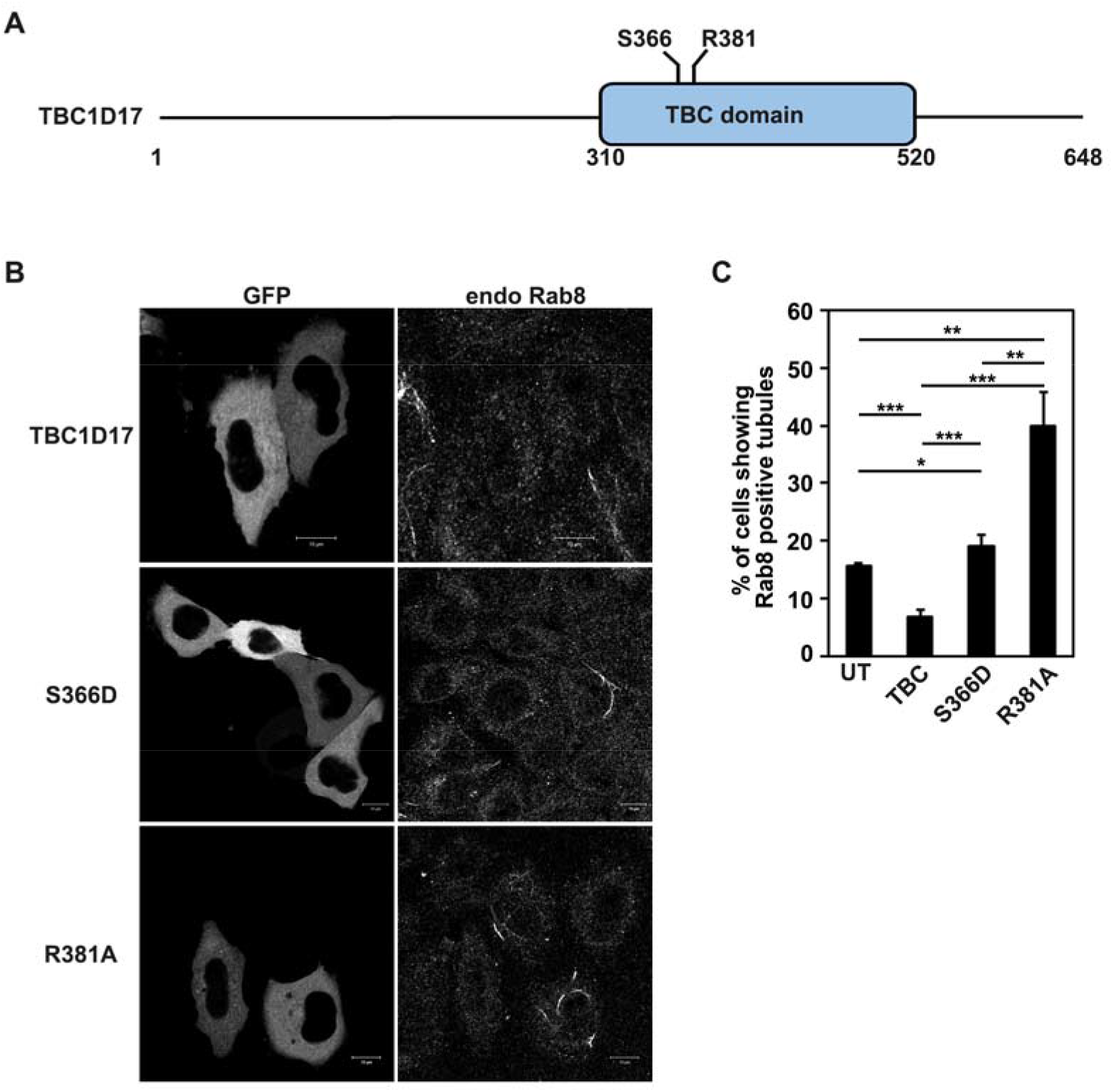
A phospho-mimicking mutant of TBC1D17 is unable to inhibit RAB8-positive tubules. (A) Schematic of TBC1D17 protein. (B) Hela cells were transfected with GFP-tagged TBC1D17 or its phospho-mimicking S366D mutant or catalytically inactive R381A mutant, and after 24 hours the cells were fixed and stained with anti-RAB8 antibody. The cells were then observed by confocal microscopy for RAB8-positive tubules. Scale bar: 10 μm. (C) Graph shows percentage of cells exhibiting RAB8-positive tubules in each scoring category. Data from three separate experiments are shown as the mean±s.d., ***P<0.0001, **P<0.01, *P<0.05.

### Possible role of phosphorylation of TBC1D17 in the transferrin-induced formation of RAB8-positive tubules

Treatment of cells with transferrin leads to activation of RAB8, as evidenced by more cells showing RAB8-positive tubules; this may be a result of inactivation of its GAP, TBC1D17. Because AKT is required for transferrin-induced formation of RAB8-positive tubules, we hypothesized that phosphorylation of TBC1D17 by AKT directly (or indirectly through some other kinase) may lead to its inactivation. Therefore, we did a bioinformatics analysis of TBC1D17 protein to look for possible sites of phosphorylation by AKT and other kinases. The consensus sequence for phosphorylation by AKT is RXRXXS/TF/L. One of the sites, S366 (EQE**R**RN**S**^**366**^**L**LHGYPRS) showed a good score, and hence an appropriate site for creating phospho-mimicking mutant by site-directed mutagenesis. S366 is located in the catalytic TBC domain of TBC1D17 (Fig. 7A). TBC1D17 and its phospho-mimicking S366D mutants were expressed in HeLa cells and RAB8-positive tubules were looked for by confocal microscopy. Unlike wild-type TBC1D17, the S366D mutant did not show reduction in the percentage of cells showing RAB8-positive tubules, indicating it to be an inactive form (Fig. 7B, C). However, this inactivity was not comparable to R381A-TBC1D17, which showed increased percentage of cells with RAB8-positive tubules (Fig. 7B, C). These results suggest that phosphorylation of TBC1D17 at S366 may be one of the mechanisms by which it might get inactivated by AKT.

### SRC kinase and PI3-kinase are required for transferrin-induced formation of RAB8 and MICAL-L1 tubules

We explored the role of SRC kinase and PI3-kinase in transferrin-induced formation of RAB8 and MICAL-L1-positive tubules, because these kinases are known to be activated by transferrin treatment of cells [3, 30]. HeLa cells grown on coverslips were serum starved for 2 hours, treated with 10μM of the SRC-inhibitor SU6646 or 250nM of the PI3-kinase inhibitor, Wortmannin along with 10μg/ml of transferrin or BSA for 1hr at 4°C followed by incubation at 37°C for 20 minutes. Cells were fixed and stained for endogenous RAB8 and MICAL-L1, and observed by confocal microscope. Representative confocal images are shown in Fig. 8A. Treatment with SRC inhibitor resulted in nearly complete inhibition of transferrin-induced formation of RAB8 and MICAL-L1-positive tubules (Fig. 8A, B, C). Treatment with PI3-kinase inhibitor also resulted in reduction of transferrin-induced formation of RAB8 and MICAL-L1-positive tubules (Fig. 8A, B, C).

**Figure 8:**
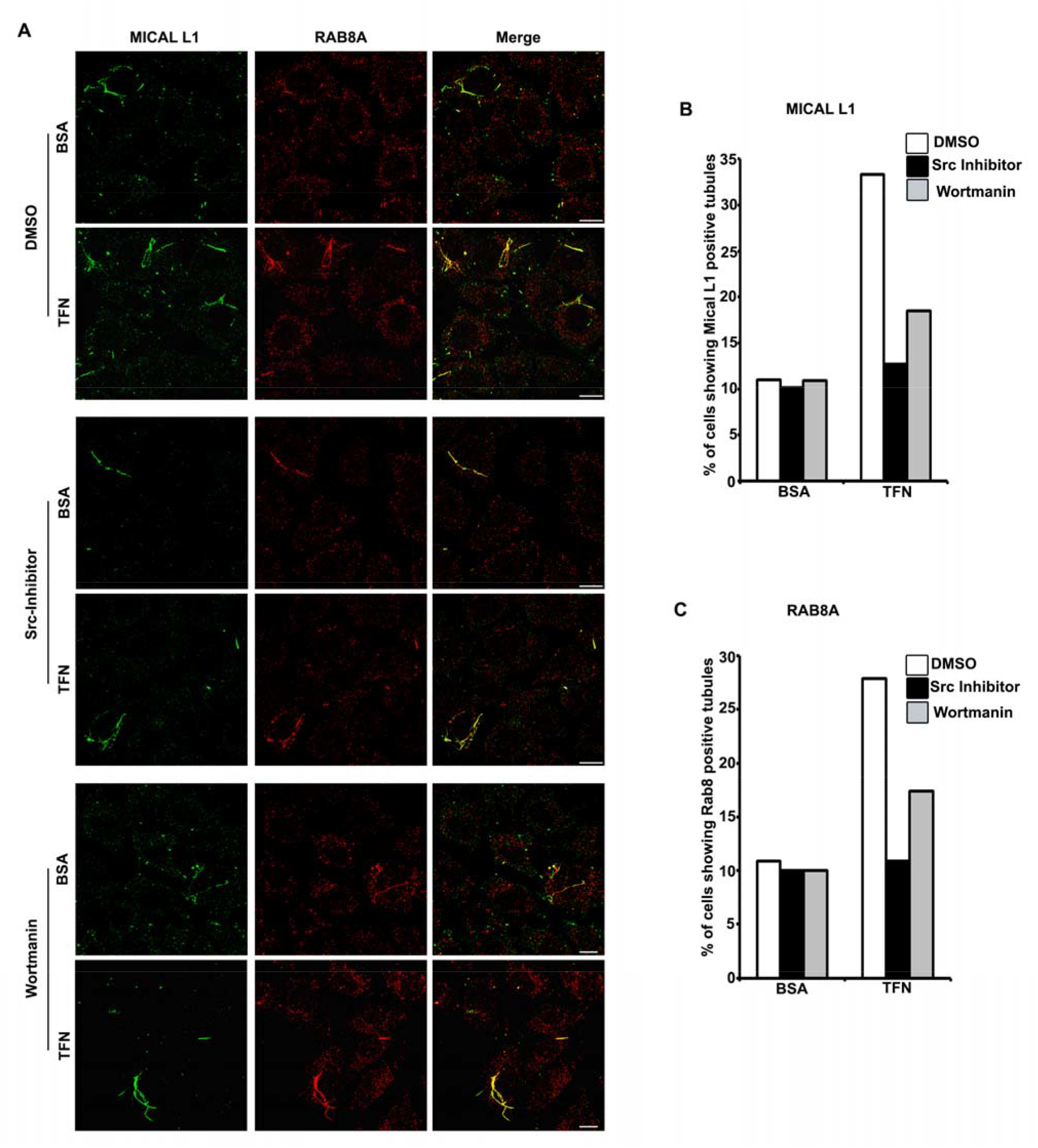
Src-kinase and PI3-kinase are required for transferrin induced formation of RAB 8 and MICAL-L1 tubules. HeLa cells grown on coverslips were serum starved for 2 hours, treated with BSA or transferrin as well as with DMSO (0.1%) or SU6656(Src kinase inhibitor10uM) or Wortmanin (250nM), kept at 4°C for 1 hour followed by incubation at 37°C for 20 minutes. Cells were fixed and stained for endogenous RAB8 and MICAL-L1 and observed by confocal microscope. Representative confocal images are shown (A) Scale bar: 10 μm. Bar diagrams show percentage of cells exhibiting MICAL-L1-positive (B) or RAB8-positive (C) tubules in each scoring category. Data from two separate experiments done with 100-165 cells are shown.

### SRC kinase inhibitor abrogates transferrin-induced AKT activation

Treatment of serum starved HeLa cells with transferrin resulted in activation of AKT as seen by phosphorylation of AKT by using a phospho-AKT (pSer473) specific antibody (Fig. 9A). Treatment of cells with SRC family kinase inhibitor SU6656 resulted in abrogation of transferrin induced increase in AKT phosphorylation (Fig. 9B). These results along with results described earlier provide evidence for the hypothesis that transferrin-induced activation of AKT mediated by SRC (or a SRC family kinase) is involved in RAB8 tubule formation.

**Figure 9:**
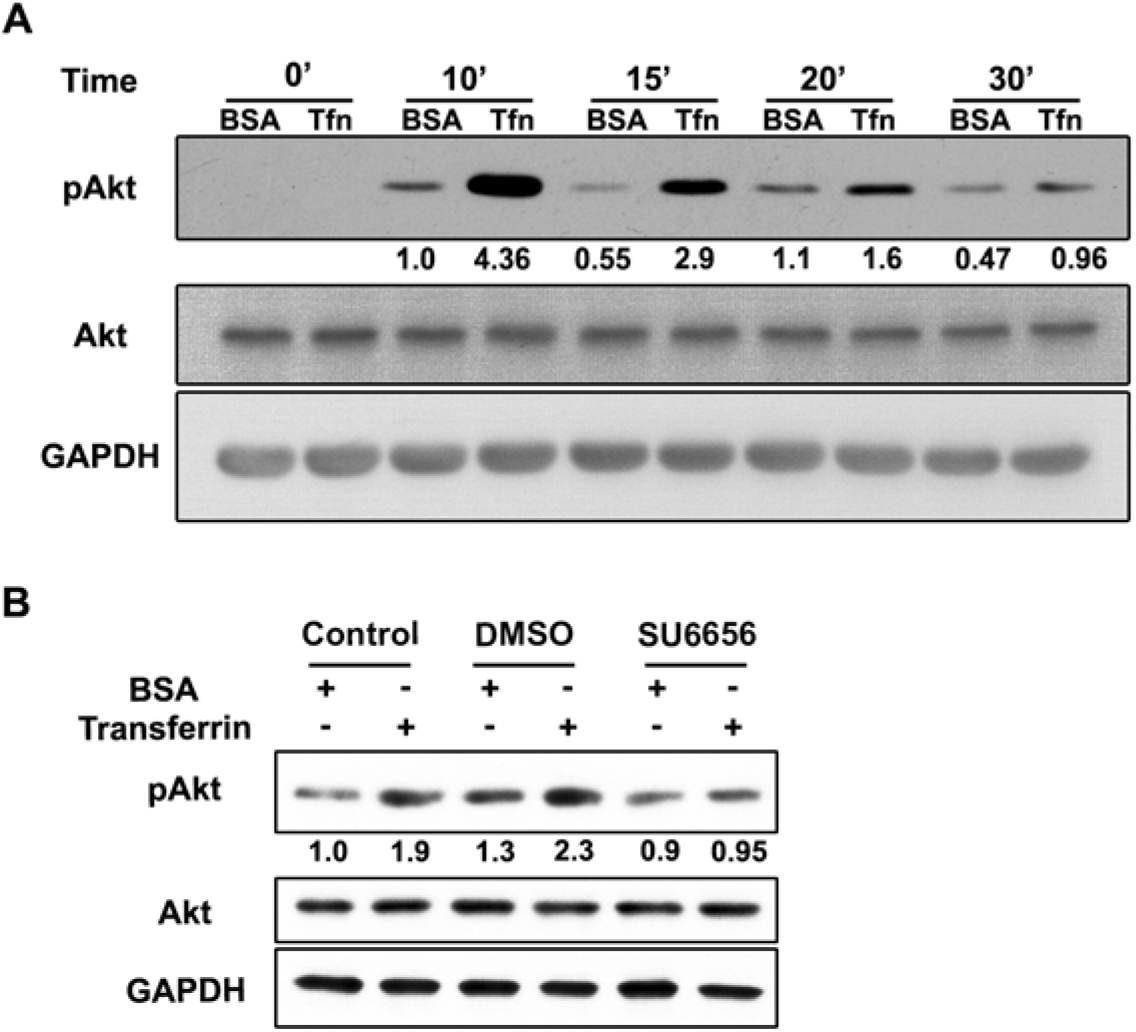
SRC inhibitor abrogates transferrin-induced AKT activation. (A) HeLa cells were serum starved for 2 hours, treated with BSA or transferrin, kept at 4°C for 1 hour followed by incubation at 37°C for 0-30 minutes. Subsequently, cell lysates were prepared and analyzed by western blotting using phospho-AKT (p-Ser473) (pAKT), AKT and GAPDH antibodies. (B) HeLa cells were serum starved for 2 hours, treated with BSA or transferrin as well as with DMSO (0.1%) or SU6656 (Src kinase inhibitor, 10 μM), kept at 4°C for 1 hour followed by incubation at 37°C for 20 minutes. Cell lysates were subjected to western blotting using indicated antibodies.

## Discussion

Binding of transferrin to its receptor initiates endocytosis of Tfn-TFRC complex leading to formation of endocytic vesicles, which move to early endosome. Iron is released in the endosomes and Tfn-TFRC complex is recycled back to the plasma membrane or moves to ERC and then recycled back to the plasma membrane. Tubules emanating from the ERC play a crucial role in recycling of TFRC and some other receptors to plasma membrane. Since endocytic trafficking and recycling of Tfn-TFRC is initiated by binding of Tfn to TFRC, we hypothesized that Tfn-induced signaling mediated by TFRC may be involved in regulating the formation of RAB8 and MICAL-L1-positive tubules. In this study, we have shown that Tfn treatment of cells induces signaling mediated by TFRC, that leads to transient increase in the formation of RAB8 and MICAL-L1-positive tubules, which requires endocytosis. This transferrin-induced increase in formation of RAB8 and MICAL-L1-positive tubules is mediated by AKT kinase, as seen by the inhibitory effect of a chemical inhibitor of AKT as well as a kinase dead AKT mutant on transferrin-induced increase in the number of cells showing RAB8 and MICAL-L1-positive tubules. Our results show that AKT activity is required not only for the formation of RAB8 and MICAL-L1-positive tubules, but also for endocytic recycling of transferrin.

### Role of AKT in regulating RAB8 activity through a GAP protein

How does AKT kinase activity regulate the formation of RAB8-positive tubules? Only the activated form of RAB8 is present on the tubules and the activity of RAB8 GTPase is negatively regulated by a GAP, TBC1D17, which also inhibits TFRC recycling [15]. Bioinformatics analysis showed several potential phosphorylation sites on TBC1D17 and Ser366 was predicted to be a potential site for phosphorylation by AKT (and some other kinases). A phospho-mimicking mutant of this site, S366D was unable to inhibit RAB8 tubule formation in normally growing cells. This indicates that phosphorylation of Ser366 of TBC1D17 by AKT might be involved in negatively regulating GAP activity of TBC1D17. A catalytically inactive mutant, R381A of TBC1D17 did not inhibit RAB8 or MICAL-L1-positive tubule formation. In fact, R381A mutant increased the percentage of cells showing RAB8-positive tubules. This indicates that S366D mutant may not be completely inactive, and phosphorylation at additional sites may be required for full inactivation of TBC1D17 by AKT directly or indirectly. This may be one of the mechanisms by which AKT regulates formation of RAB8 tubules. This is shown schematically in Fig. 10.

**Figure 10.**
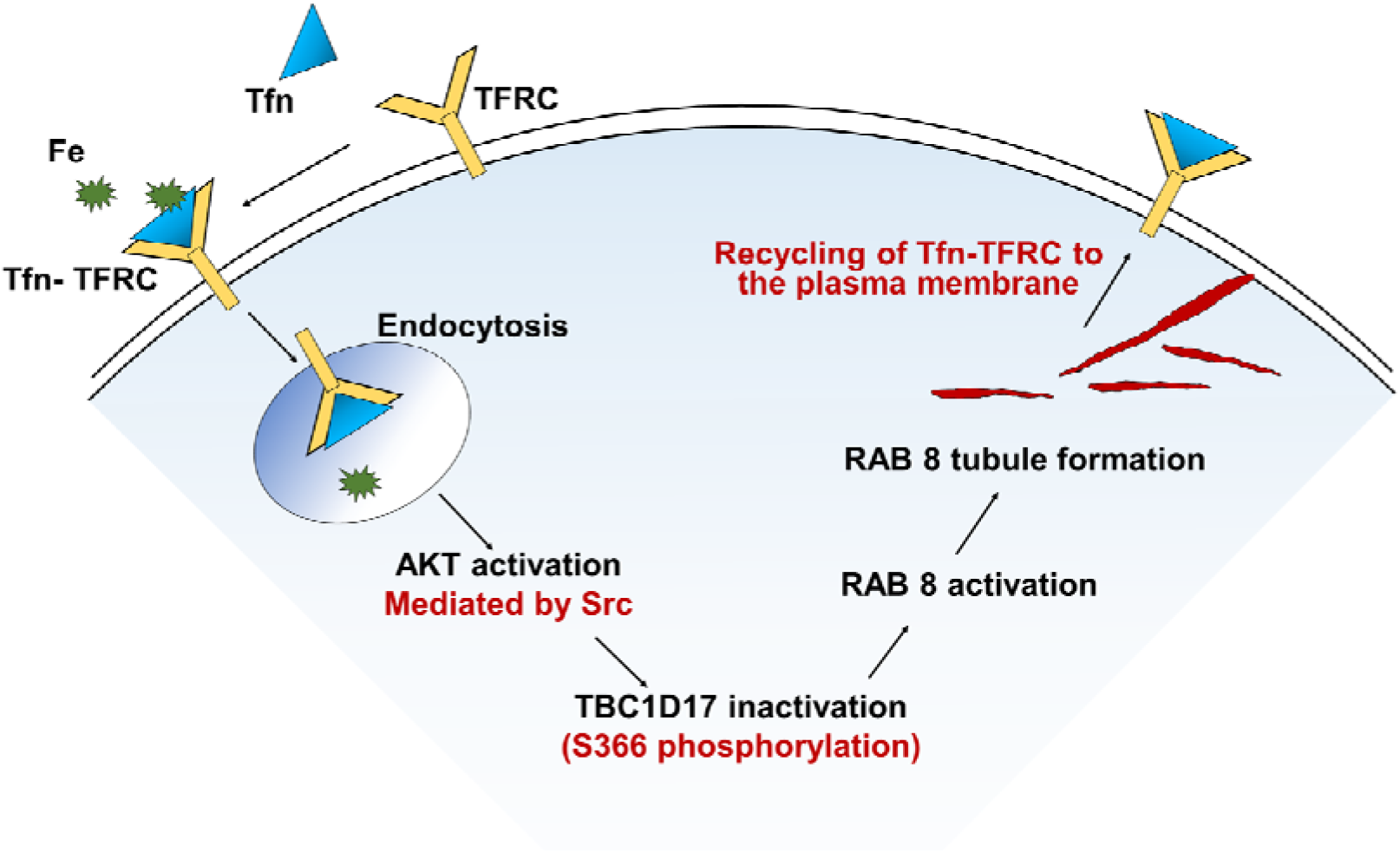
Schematic showing proposed role of AKT kinase in transferrin-induced formation of RAB8-positive tubules involved in recycling of TFRC. Binding of transferrin to its receptor TFRC, a transmembrane protein leads to endocytosis. Activation of SRC upon binding of transferrin to its receptor has been reported earlier [30]. Activated SRC leads to AKT activation, which then inactivates TBC1D17 (a GAP for RAB8) possibly through phosphorylation at S366. This results in RAB8 activation which is recruited to the tubules. Activated RAB8 facilitates recruitment of MICAL-L1 to the tubules. These tubules mediate recycling of transferrin-TFRC complex to the plasma membrane.

Previously, regulation of catalytic activity of GAP proteins by AKT mediated phosphorylation has been reported. Insulin stimulated increase in glucose uptake is mediated by glucose transporter GLUT4, which is translocated from the storage vesicles to the plasma membrane. This GLUT4 translocation is regulated by AKT mediated phosphorylation of GAP proteins, TBC1D4 (also known as AS160) and TBC1D1 [36-39]. These TBC proteins, TBC1D4 and TBC1D1 regulate several RABs, particularly RAB10 [39]. Regulation of TBC1D17 by AMPK-dependent phosphorylation has been reported in the control of RAB5 activity and glucose uptake [40]. Thus, regulation of activity of TBC proteins by phosphorylation is an important mechanism to control the functions of RAB GTPases.

### What is the function of slow recycling of TFRC?

The main function of transferrin and TFRC in the cells is uptake of iron. Iron is released from transferrin in the early endosomes, from where it is transported into the cytosol by iron transporters. Transferrin-TFRC complex is transported back to plasma membrane through two pathways: fast recycling from early endosomes, and slow recycling from recycling endosomes. What is the purpose of slow recycling of TFRC? Fast recycling of TFRC should be sufficient for iron uptake. This implies that slow recycling must serve some other function. One possibility is that signal transduction by TFRC, which sometimes requires endocytosis, may require its trafficking to recycling compartment. But it is not clear whether trafficking of TFRC beyond early endosomes is required for signaling function of TFRC. Another possibility is that TFRC delivers membrane to recycling endosomes and autophagosomes. Delivery of membrane from recycling endosomes to autophagosomes for their biogenesis has been shown earlier [41, 42]. Recently, it has been shown that autophagy receptor optineurin (OPTN) plays a crucial role in the delivery of membrane from TFRC-positive endosomes or recycling endosomes to autophagosomes [43, 44]. Whether membrane delivery to autophagosomes occurs from TFRC-positive early endosomes or recycling endosomes (or both) is not clear. However, it is interesting to note that a glaucoma-associated mutant M98K-OPTN shows enhanced delivery of TFRC and possibly associated membrane to autophagosomes in retinal cells [8]. Another mutant of OPTN, E478G, associated with amyotrophic lateral sclerosis, is defective in trafficking of TFRC and possibly associated membrane to autophagosomes [43].

### The relationship between RAB8 and MICAL-L1-positive tubules

The relationship between RAB8 and MICAL-L1-positive tubules is somewhat complex [18]. We observed that the increased formation of RAB8 and MICAL-L1-positive tubules in response to Tfn treatment follows similar time course. Inhibitors of endocytosis, SRC kinase and AKT kinase showed similar effect on the Tfn-induced formation of RAB8 and MICAL-L1-positive tubules. Furthermore, RAB8 and MICAL-L1 showed good colocalization on the tubules but not elsewhere in the cells. TBC1D17, a GAP for RAB8 inhibits formation of RAB8 as well as MICAL-L1 tubules. These results are consistent with the suggestion that activated RAB8 facilitates recruitment of MICAL-L1 on the tubules. However, we noticed that R381A mutant of TBC1D17 enhanced the formation of RAB8-positive tubules but had no effect on MICAL-L1-positive tubules. This indicates that there are additional mechanisms for regulating the formation of MICAL-L1-positive tubules.

In conclusion, our results show that binding of transferrin to TFRC induces AKT mediated signaling that regulates formation of RAB8 and MICAL-L1-positive tubules involved in recycling of transferrin receptor. This signaling is dependent on endocytosis. Close association of MICAL-L1-positive tubules with the formation of RAB8-positive tubules under various conditions indicates that activated RAB8 facilitates recruitment of MICAL-L1 to the tubules.

## Acknowledgements

This work was supported by a grant [J.C. Bose National Fellowship (SR/S2/JCB-41/2010)] awarded to G.S from the Science and Engineering Research Board, Department of Science and Technology, Government of India. G.S. acknowledges Indian National Science Academy for a Senior Scientist grant.

## Notes

### Competing Interest Statement

The authors have declared no competing interest.

